# CancerInSilico: An R/Bioconductor package for combining mathematical and statistical modeling to simulate time course bulk and single cell gene expression data in cancer

**DOI:** 10.1101/328807

**Authors:** Thomas D Sherman, Luciane T Kagohara, Raymon Cao, Raymond Cheng, Matthew Satriano, Michael Considine, Gabriel Krigsfeld, Ruchira Ranaweera, Yong Tang, Sandra A Jablonski, Genevieve Stein-O’Brien, Daria A Gaykalova, Louis M Weiner, Christine H Chung, Elana J Fertig

**Affiliations:** Department of Oncology, Division of Biostatistics and Bioinformatics, Sidney Kimmel Comprehensive Cancer Center, Johns Hopkins University, Baltimore, MD USA; Science, Math and Computer Science Magnet Program, Poolesville High School, Poolesville, MD USA; Department of Mathematics, University of Waterloo, Waterloo, Ontario, Canada; Moffitt Cancer Center, Tampa, FL, USA; Salubris Biotherapeutics, Inc, Gaithersburg, MD, USA; Lombardi Comprehensive Cancer Center, Georgetown University, Washington, DC USA; Institute of Genetic Medicine, Johns Hopkins University, Baltimore, MD USA; Department of Otolaryngology-Head and Neck Surgery, Johns Hopkins University School of Medicine, Baltimore, MD USA; Department of Applied Mathematics and Statistics, Johns Hopkins University; Department of Biomedical Engineering, Johns Hopkins University

## Abstract

Bioinformatics techniques to analyze time course bulk and single cell omics data are advancing. The absence of a known ground truth of the dynamics of molecular changes challenges benchmarking their performance on real data. Realistic simulated time-course datasets are essential to assess the performance of time course bioinformatics algorithms. We develop an R/Bioconductor package, *CancerInSilico*, to simulate bulk and single cell transcriptional data from a known ground truth obtained from mathematical models of cellular systems. This package contains a general R infrastructure for running cell-based models and simulating gene expression data based on the model states. We show how to use this package to simulate a gene expression data set and consequently benchmark analysis methods on this data set with a known ground truth. The package is freely available via Bioconductor: http://bioconductor.org/packages/CancerInSilico/

## Introduction

Time course bioinformatics analysis techniques are emerging to delineate cellular composition and pathway activation from longitudinal genomics data [1,2]. However, benchmarking their performance is challenged by a lack of ground truth of the processes occurring in those datasets. For example, even relatively simple covariates, such as cellular density and proliferation rates impact experimental measures at a given time point, such as therapeutic sensitivity in cancer [3]. The interactions between these processes will introduce additional correlation structure between genes measured with genomics technologies. Simulated data can enable robust benchmarking of bioinformatics analysis methods for omics data. Statistical methods that utilize expected gene expression profiles from reference datasets to model the error distribution of bulk and single cell sequencing data are prominent [4–6]. Yet, there are few time course omics datasets to use as a benchmark and even fewer with known cellular-molecular dynamics. Therefore, new simulation systems with known ground truth are needed to benchmark the performance of emerging time course bioinformatics algorithms for bulk and single cell datasets.

Mathematical models of cellular dynamics are maturing in systems biology and can be used to track the state of the processes occurring in each cell in complex biological systems, such as cancer [7–13]. Some models simulate cell growth at a cellular level, where the population behavior is driven by the laws governing the individual cells and their interactions [14,15]. To further capture the complexity of biological systems, numerous multiscale and hybrid models linking cellular signaling to the equations of the cellular composition are emerging [16–18]. These models often require numerous parameters to simulate high throughput proteomic and transcriptional data and therefore often have similar complexity to real biological systems. Thus, mathematical models provide a robust framework from which to develop simulated time course datasets that are reflective of biological systems.

In this paper, we present a new software package to simulate time course transcriptional data. This is done by developing a general software framework to integrate mathematical models of cellular growth with statistical models of genomics data. The software is implemented in the R/Bioconductor package *CancerInSilico*. We simulate pathway activity based upon the simulated distribution of growth factor, state in the cell cycle, and cellular type. We couple a mathematical model from [14] with a statistical model from [19] to simulate transcriptional data based upon simulated pathway activity. We simulate data from microarrays and single cell RNA-seq using established platform-specific error distribution models [4,19,22]. Finally, we demonstrate how this framework can be used to benchmark time course analysis tools for genomics data.

## Design and implementation

### Software architecture

*CancerInSilico* is designed with an R user interface so that it is familiar to the bioinformatics community. The components of the simulation such as cell types and pathways are implemented as S4 classes in R. The cell model component of the simulation is implemented as an S4 class hierarchy, where features such as cell geometry (on-lattice vs off-lattice) form the basis for a set of models that individual implementations can inherit from. This allows different levels of the cell model to be considered separately so that the user can examine the effects of not just a single model but a whole class of models. The hierarchy also simplifies the number of parameters the user must interact with. Each level of the hierarchy contains its share of the overall parameters, so if the user wants to modify the low level implementation parameters, they can do so without worrying about any effects to the parameters upstream. This object oriented design simplifies the workflow by allowing each component of the model to be specified separately. In order to run the simulation, the user just needs to pass in any desired components along with a few high-level parameters.

All components of the simulation, including the cell model hierarchy, have a mirrored class structure in C++. While the classes in R contain the necessary parameters, the C++ classes also include the necessary routines to efficiently simulate a cell model. This architecture essentially allows the user to see a snapshot of the model in R and trust that the C++ backend will run it exactly as they prescribe. Moreover, this combines a simple user interface in R with a powerful, efficient backend in C++. The C++ library is exposed to R using the Rcpp package from CRAN.

The statistical model for gene expression simulation is written in R and exists outside of the previously mentioned class structure. It is intended to be an independent component that only needs the output of a cell simulation and a set of parameters in order to run. This way it is agnostic to any implementation details of the cell simulation, as well as any components that may be added to the cell simulation. The needed parameters are provided as an S4 class for convenience, allowing them to easily be saved alongside the simulation results.

### Implementation details

The cellular growth simulation is driven by the cell model hierarchy and the peripheral components that can be added such as drugs and cell types. The `CellBasedModel` class at the top of the hierarchy specifies the relationship between a cell model and any peripheral components. It makes no requirements on the cell geometry or updating procedure of the model. In most cases, the user will be modifying parameters at this level. The model-component relationship is implemented with virtual functions in C++ so individual model implementations can override the default behavior. This top level class also uses pure virtual functions to specify the functions a cell model must implement. We note that the `CellBasedModel` has an associated `Cell` class to separate cell-level logic with model-level logic. This is a design pattern seen across all levels of the hierarchy.

Any class that fully implements the specification of a `CellBasedModel` can be used in *CancerInSilico*, however it is convenient to define an intermediate class between the top-level `CellBasedModel` and an actual implementation. This layer describes a certain set of cell models, usually by specifying cell geometry, e.g. off-lattice vs on-lattice. This is a useful abstraction since the most computationally expensive aspects of a cell model often stem from the cell population data structure. If every model implementation required designing this data structure from scratch it would put an enormous burden on the developer. By having a pre-defined, efficient data storage and access API, new cell dynamics can be quickly prototyped and will come with an expected level of performance.

The intermediate layer specifies the cell geometry and structure, but the updating procedure must be handled by the actual cell model implementation. This lowest-level of the hierarchy is responsible for actually enforcing the desired cell mechanics. This is typically done by specifying some total energy function on the full cell population. Updating then involves several kinds of changes that are either accepted or rejected with a probability based on the total energy function. Most cell models in the literature [12, 14] can be identified by how they handle this updating procedure. By isolating this layer within the *CancerInSilico* architecture, such models can be easily implemented.

While the cellular growth simulation is an important part of *CancerInSilico*, the main feature of the package is the gene expression simulation. The connection between this simulation and the cellular growth simulation happens through user defined pathways that are then associated with gene expression changes in a corresponding set of genes. We define an intermediate variable (*P*) that is a continuous value between zero and one that records how active each biological pathway is within each modeled cell. The value of *P* in a given cell at a given time is determined by a user defined function in R. This allows for many types of pathways to be considered and for a great deal of expressiveness in how each pathway behaves. *CancerInSilico* also comes with some pre-defined pathways so that the burden is not entirely on the user to design the pathway behavior. Along with this user defined function, each pathway also has a set of genes annotated to it. Each gene in this set has a pre-specified expression range (*Gmin* to *Gmax*), determined either from a reference dataset or according to a specified distribution.

The function `inSilicoGeneExpression` combines the results of a call to `inSilicoCellModel` and a list of user defined pathways to simulate the requested type of transcriptional data. The first step is to create a matrix of mean expression values. This is done by evaluating each gene according to the specified behavior from the list of pathways. The expected expression value for each gene and each sample is given as:

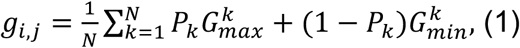

where i indexes each gene, j each sample, and k each pathway. We note that while the mean used in eq (1) is provided as the default, *CancerInSilico* allows for user defined functions to combine pathway specific expression values. In bulk data, *P_k_* is determined by computing the average value for pathway activity in a random set of *N* sampled cells, whereas in single cell data the value of *P_k_* for each of the *N* sampled cells is used directly. The simulated gene expression value is obtained using a platform specific measurement error based on this expectation. A normal error model is used to simulate log transformed microarray data and a negative binomial error model, adapted from the code for LIMMA voom [22], is used to simulate bulk RNA-sequencing data. Measurement error for simulated single cell RNA-sequencing data are generated using the error and drop out models from Splatter [4].

## Results

### The *CancerInSilico* workflow

#### Running a cell simulation with *inSilicoCellModel*

The first step when simulating gene expression data with *CancerInSilico* is to create a cell simulation to serve as a reference point. This cell simulation will represent the underlying cellular processes driving the gene expression profiles. In order to run a cell simulation, we must call `inSilicoCellModel`. The three required arguments to this function are the initial number of cells in the simulation, the number of hours to simulate, and the initial density of the cell population. Optionally, it is possible to select the underlying mathematical model. A full description of the optional parameters is included in the supplemental material. An example call to the function might look like:

~~~
> cellModel = inSilicoCellModel(100, 72, 0.01, “DrasdoHohme”)
~~~

#### Defining pathways

Before we can move on to simulating gene expression data, it is necessary to define the pathways which link the cell model state to the activity among a set of genes. *CancerInSilico* comes with a set of default pathways, however we can also explicitly define new pathways. In order to create a new pathway we must specify the names of the genes in the pathway and activity function which takes a cell model as an argument and returns a value between zero and one based on how active the pathway is at the current time point. Once a pathway is defined it must be calibrated either to a real data set or using a statistical distribution. This calibration step is important so that the range of the gene expression values is reasonable. Here is an example of calibrating a default pathway with a distribution. The mean expression levels for all genes is exponentially distributed and the range of expression values per gene is normally distributed.

~~~
> data(samplePathways) # load pwyMitosis, pwySPhase
> pwyMitosis = calibratePathway(pwyMitosis, lambda=20, stddev=2)
> pwySPhase = calibratePathway(pwySPhase, lambda=20, stddev=2)
> pathways = list(pwyMitosis, pwySPhase)
~~~

#### Simulating gene expression data with *inSilicoGeneExpression*

Now that we have a defined set of pathways and a completed cell simulation, we are able to simulate a gene expression data set. This step involves computing the mean level of expression in all the pathway genes and applying a statistical error model based on the type of data being generated. There are a few parameters that control this part of the simulation. `sampleFreq` and `nCells` specify how often samples are drawn and how many cells are in each sample. `RNAseq` and `singleCell` are Boolean parameters that specify the type of data to generate. A full description of the parameters can be found in the supplemental material. An example call to the function might look like:

~~~
> params = new(“GeneExpressionParams”, nCells=50, RNAseq=TRUE)
> exp = inSilicoGeneExpression(cellModel, pathways, params)
~~~

### Example: Simulating time-course bulk data

Using the workflow described in the previous section, we simulate a microarray data set across 43 time points. The underlying mathematical model for cellular growth in this case is an off-lattice, cell-center model from Drasdo and Höhme [14]. We model pathways related to the phase transition from G to S and G to M, as well as a pathway related to contact inhibition. For the G to M and G to S pathways, the pathway activity is either zero or one at the current time point depending on whether or not the cell is transitioning phases. The contact inhibition pathway activity is defined by the “local density” of the cell, which is the proportion of surrounding area of a cell that is occupied by other cells. We run the cell model for 168 hours, enough for the cell population growth to slow down due to the density of the cells, and simulate gene expression for 150 genes. We generate a heatmap of the data using the `heatmap.2` function in R **(Figure 2a)**. We also run PCA on the resulting microarray data set and show that, as expected, time and cell phase are the processes driving the simulated gene expression **(Figure 2bc)**.

**Fig 1.**
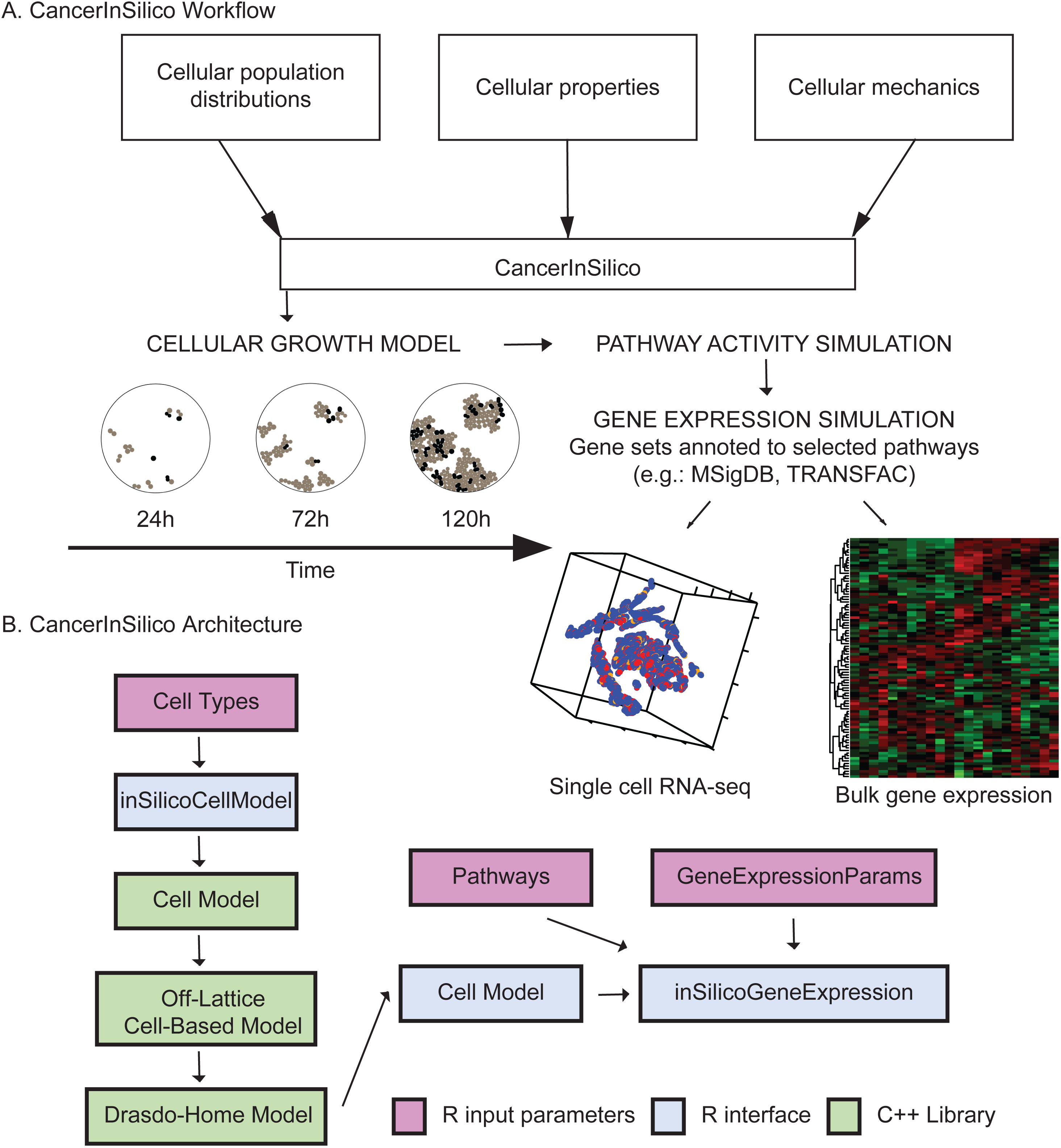
Visual representation of CancerInSilico. (a) Overview of association between mathematical and statistical modeling to generate simulated time course gene expression data. (b) Each component of the software is shown with arrows indicating how it is related to the rest of the components. R input arguments shown in pink, R software components shown in blue, and C++ components shown in green.

**Fig 2.**
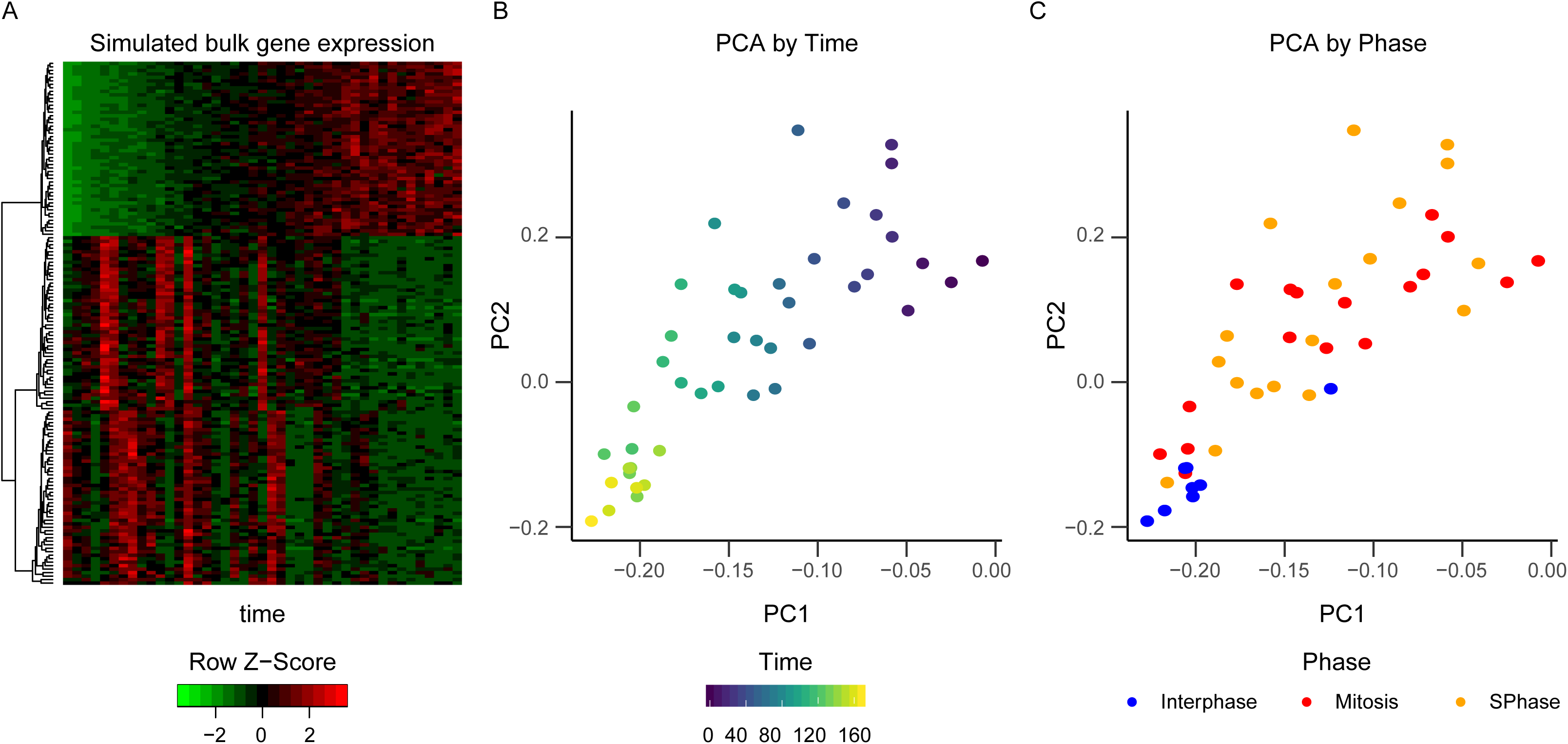
PCA of simulated time course microarray data. (a) Heatmap of microarray data. Plot of first two principal components colored by (b) time and (c) cell phase.

### Example: Simulating time-course single-cell data

*CancerInSilico* also encodes an option to simulate single cell RNA-sequencing data to generate omics data that reflects the heterogeneity of the sample population. To model this heterogeneity, *CancerInSilico* allows us to label each cell as being from a distinct cell type. We have control over the distribution of cell-cycle lengths within each cell type through a user defined function in R. We apply this framework to model two distinct cell types, one with a mean cell cycle length of 12 hours and standard deviation of 4 hours (type A) and one with a mean cell cycle length of 36 hours and standard deviation of 4 hours (type B). This simulation models the pathway activity and corresponding gene expression changes for each cell with a negative binomial error model and dropout model adapted from Splatter [4]. The model then randomly samples a pre-specified number of cells. We apply this technique to simulate single cell RNA-sequencing data from a simulation of a population equally distributed between the two types described above (**Fig 3**). Each cell type is labeled as a pathway with binary values for activity to activate a gene set that corresponds to cellular identity. In this simulated single-cell RNA-seq data, we observe strong separation between cell types (**Fig 3a**) and time (**Fig 3b**) and observe a mixture between cell cycle phases (**Fig 3c**).

**Fig 3.**
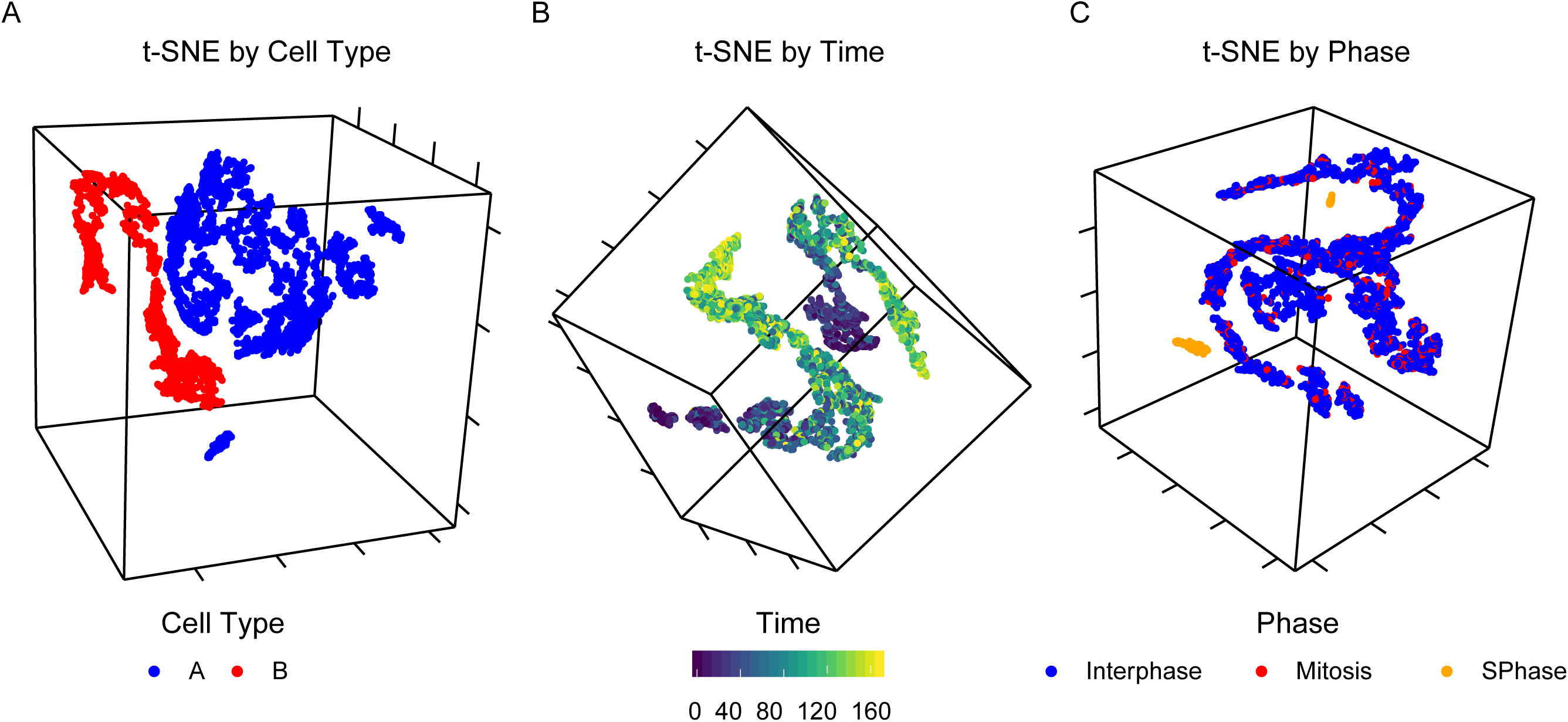
T-SNE of simulated time course single cell RNA-sequencing data for a population with cells of types A and B. Points colored by (a) cell type, (b) time, and (c) cell cycle phase.

### Example: Benchmarking analysis tools

We can use the wide range of simulation conditions in *CancerInSilico* to benchmark time course gene expression analysis methods. Moreover, by starting with biological parameters as opposed to statistical parameters, we can benchmark analysis methods on conditions we actually care about. *CancerInSilico* is particularly useful when the cellular processes underlying the simulation of interest have complex relationships with each other. This complexity is handled by the underlying cellular growth simulation, and while it does not perfectly capture the dependence between cellular processes, it does provide a standardized, justifiable method. Furthermore, *CancerInSilico* allows for the underlying cellular growth model to be easily swapped out so the benchmark itself can be tested for sensitivity to a particular model.

Here we explore the effects of dependent cellular processes when using Independent Component Analysis (ICA), implemented in the R package `fastICA`, to analyze our simulated RNA-seq data set. We define two cell-types which have identical properties and provide an associated pathway for each one. We also define a third pathway which is proportional to the growth rate of a cell. We have two datasets, one in which all pathways have a distinct set of genes, and one where the growth pathway contains all of the genes in each cell type specific pathway. In this way, the growth pathway is confounding the cell type specific pathways. We can see that ICA with two components perfectly separates the cells (**Fig 4a**) and temporal dynamics (**Fig 4b**) of the simulated data with no confounding. When the overlapping pathway is introduced some of the cells can no longer be separated by type (**Fig 4c**) although the temporal dynamics are still separable via ICA (**Fig 4d**). Thus, this provides one example of the utility of *CancerInSilico* to benchmark the sensitivity of a time course analysis algorithm to its underlying mathematical assumptions. Additional parameters may be varied and further cell cycle pathways introduced to the data to increase the complexity of these simulations and more closely mirror the complexity of the analysis tasks in real, time course genomics data.

**Fig 4.**
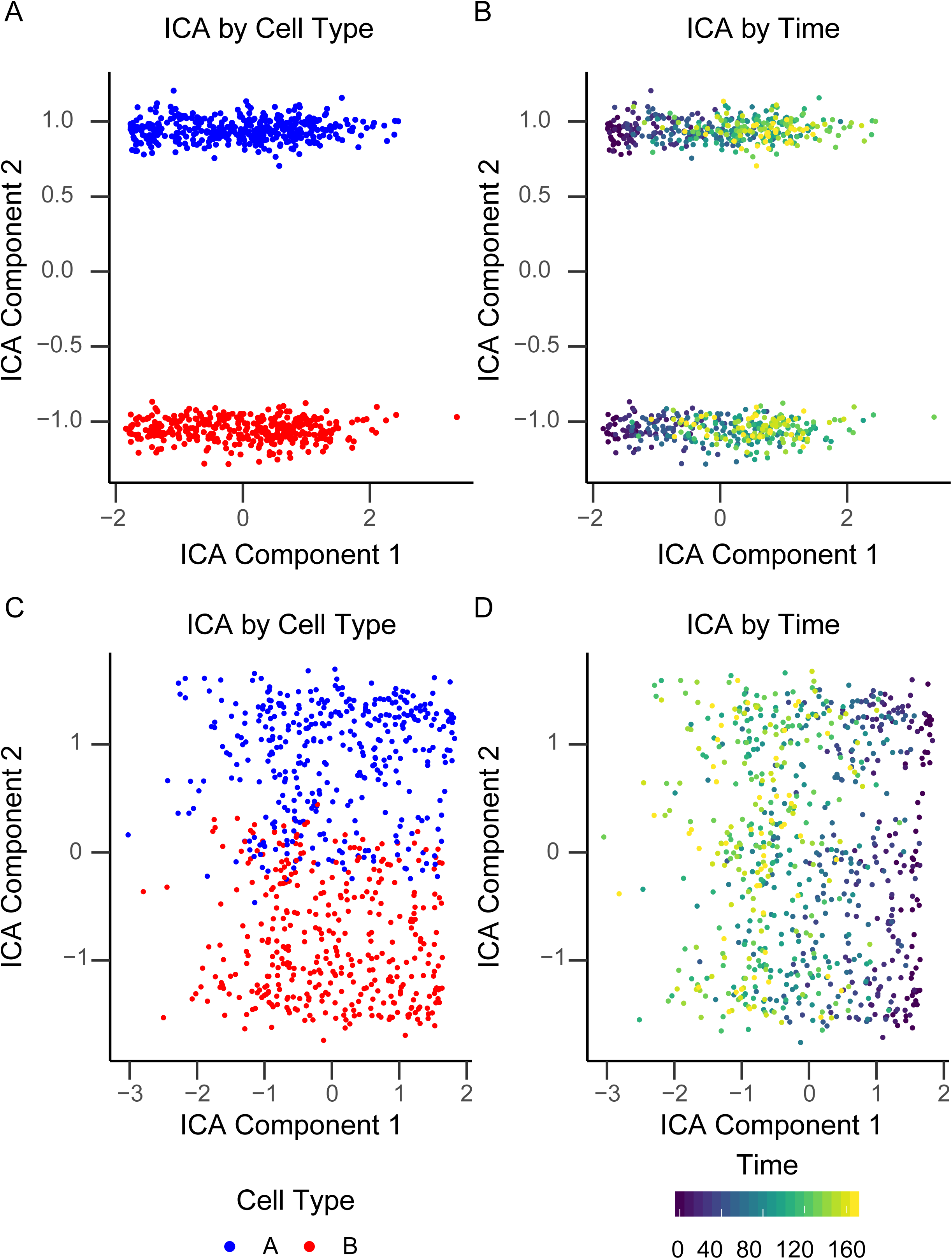
Benchmarking ICA when cell type pathways are confounded with third pathway. ICA with no confounding colored by (a) cell type and (b) time. ICA with third pathway confounding colored by (c) cell type and (d) time.

### Availability and future directions

We develop a new R/Bioconductor package *CancerInSilico* that couples mathematical models of cellular growth with statistical models of technical noise. Using this coupling to model changes in gene sets annotated to cell signaling pathways [4,19,22] enables simulation of time course bulk omics data. The modeling of individual cells in this system also enables simulation of time course, single cell RNA-sequencing data. *CancerInSilico* provides a wide range of parameter spaces for the user to explore when simulating time-course gene expression data and the modular design makes it possible to swap different cell models in and out of an existing simulation. Thus, this package provides software that can be used to benchmark the performance of methods for time course bioinformatics analysis from a known ground truth that is lacking in real data. We note that, to our knowledge, this is the first software package designed to simulate time course gene expression data.

The statistical models used to simulate gene expression data from pathway activity in *CancerInSilico* mirror the process by which omics tools estimate that activity. Namely, pathways are assumed to activate discrete sets of genes annotated to a common function based upon the modeled cellular state. Default parameters for the model yield omics profiles with strong separation between signaling pathways that are greatly simplified relative to those observed in real data. Therefore, applying omics algorithms to these default simulations may result in under estimation of the accuracy of their performance for real time course data. Future work is essential to benchmark the performance of the CancerInSilico algorithm to real data. Mathematical models of gene regulatory networks have been developed to model the dynamics of regulatory networks that lead to transcriptional changes [28,29]. Hybrid, multi-scale approaches that combine these network-based models with the cellular-scale models more accurately model the complexity of system-wide dynamics [16–18] and are a promising area for future work to simulate time course omics data. However, the complexity of these gene regulatory models and extensive parameterization will limit the straightforward validation of omics algorithms that is possible from the simplified statistical models employed in *CancerInSilico*. We note that the complexity of the time course data simulated with *CancerInSilico* can be tuned by modifying the overlap between genes in simulated pathways, altering cell type specific cell growth parameters, or increasing the variation in parameter values across cells. Thus, we recommend benchmarking time-course omics data analysis algorithms on simulated data generated from a wide range of these parameter values to fully assess their performance.

*CancerInSilico* is available as an R package on Bioconductor bioconductor.org/packages/CancerInSilico/ and the source code is made available at github.com/FertigLab/CancerInSilico. A live tutorial (vignette) is provided in the R package and link to a pre-rendered version is available in the GitHub README. *CancerInSilico* is supported and tested on Windows, Mac, and Linux. The source code to generate the figures seen in this paper can be found at github.com/FertigLab/CancerInSilico-Figures.

## Supporting information

## ACKNOWLEDGEMENTS

We thank Ludmila V Danilova, Alexander V Favorov, Emily Flam, Dylan Kelley, Feilim Mac Gabhann, Cristian Tomasetti, and members of NewPISlack for critical comments and feedback on this project.

