## Supplementary material for "CancerInSilico: An R/Bioconductor package for combining mathematical and statistical modeling to simulate time course bulk and single cell gene expression data in cancer"

### CancerInSilico Supplemental Material

#### Introduction

Here we make a full accounting for all parameters in the *CancerInSilico* package. There are two distinct groups of parameters, one group controls the cellular growth simulation and the other controls the gene expression simulation. The parameters for the cellular growth simulation are split up among a hierarchy of classes. In addition to the core simulation parameters, peripheral objects such as cell types and pathways can be created. For more information about such objects, see the package tutorial found here: [https://www.bioconductor.org/packages/release/bioc/vignettes/CancerInSilico/inst/doc/CancerInSilico.html](https://www.bioconductor.org/packages/release/bioc/vignettes/CancerInSilico/inst/doc/CancerInSilico.html)

All parameters are fully documented in the R package along with a description of each function. The user's manual with all documentation can be found here: [https://www.bioconductor.org/packages/release/bioc/manuals/CancerInSilico/man/CancerInSilico.pdf](https://www.bioconductor.org/packages/release/bioc/manuals/CancerInSilico/man/CancerInSilico.pdf)

#### Parameters for Cellular Growth Simulation

**CellModel**

| Parameter Name | Description | [default] Value |
|---|---|---|
| initialNum | Initial number of cells | integer > 0 |
| runTime | Total number of simulated hours | double > 0 |
| density | Initial density of the cells | 0 < double < 0.5 |
| boundary | Constrain cells within circular boundary | [1][true] boolean |
| syncCycles | Synchronize cell phase across all cells | [1][false] boolean |
| randSeed | Random seed for internal RNG | [1][0] integer |
| timeIncrement* | Amount of time passed during each time step | [inferred] double > 0 |

**OffLatticeModel**

| Parameter Name | Description | Value |
|---|---|---|
| maxTranslation* | Limits distance cells can travel in one time step | [inferred] double > 0 |
| maxRotation* | Limits angle cells can rotate in one time step | [inferred] double > 0 |

**DrasdoHohmeModel**

| Parameter Name | Description | Value |
|---|---|---|
| nG | Average number of steps between growth steps | [2][28] integer >> 1 |
| epsilon | Resistance against non-optimal update steps | [2][10] double > 0 |
| delta | Short range cell interaction distance | [2][2] 0 < double < 0.4 |

#### Parameters for Gene Expression Simulation

| Parameter Name | Description | [default] Value |
|---|---|---|
| sampleFreq | How many hours in between sample time points | [1][1] integer > 0 |
| RNAseq | Generate RNA-seq data | [1][false] boolean |
| singleCell | Generate single-cell data | [1][false] boolean |
| nCells | Number of cells to sample at each time point | [3][100] integer >> 0 |
| nDummyGenes | Number of pure noise genes | [1][0] integer |
| dummyDist | Distribution for expression of pure noise genes | R function |
| combineFUN | How to combine expression across pathways | [4][mean] R function |
| randSeed | Random seed for internal RNG | [1][0] integer |
| perError | Error for normal (microarray) model | [5][0.1] double > 0 |
| bcvCommon | Error for voom (RNA-seq) model | [5][0.2] double > 0 |
| bcvDF | Degrees of freedom for voom model | [5][40] integer > 0 |
| dropoutPresent | Simulate dropout in single cell RNA-seq | [1][false] boolean |
| dropoutMid | Parameter for dropout distribution | [6][0] double |
| dropoutShape | Parameter for dropout distribution | [6][-1] double |

* these parameters are rarely set directly, since good values can be inferred from the other parameters in the simulation

[1] Common choice for parameter – not necessarily better or worse than any other value
[2] Values found in [1]
[3] Enough cells so that averages are representative
[4] Allows for all pathways to contribute to overall activity
[5] Reasonable values based on experience – running over a range of values is recommended
[6] Found in the R package Splatter [2]
